# Genetic designs for stochastic and probabilistic biocomputing

**DOI:** 10.1101/2024.03.22.586310

**Authors:** Lewis Grozinger, Jesús Miró-Bueno, Ángel Goñi-Moreño

## Abstract

The programming of computations in living cells can be done by manipulating information flows within genetic networks. Typically, a single bit of information is encoded by a single gene’s steady state expression. Expression is discretized into high and low levels that correspond to 0 and 1 logic values, analogous to the high and low voltages in electronic logic circuits. However, the processes of molecular signaling and computation in living systems challenge this computational paradigm with their dynamic, stochastic and continuous operation. Although there is a good understanding of these phenomena in genetic networks, and there are already stochastic and probabilistic models of computation which can take on these challenges, there is currently a lack of work which puts both together to implement computations tailored to these features of living matter. Here, we design genetic networks for stochastic and probabilistic computing paradigms and develop the theory behind their operation. Moving beyond the digital abstraction, we explore the concepts of bit-streams (sequences of pulses acting as time-based signals) and probabilistic-bits or p-bits (values that can be either 1 or 0 with an assigned probability), as more suitable candidates for the encoding and processing of information in genetic networks. Specifically, the conceptualization of signals as stochastic bit-streams allows for encoding information in the frequency of random expression pulses, offering advantages such as robustness to noise. Additionally, the notion of p-bit enables the design of genetic circuits with capabilities surpassing those of current genetic logic gates, including invertibility. We design several circuits to illustrate these advantages and provide mathematical models and computational simulations that demonstrate their functionality. Our approach to stochastic and probabilistic computing in living cells not only enhances and reflects understanding of information processing in biological systems but also presents promising avenues for designing genetic circuits with advanced functionalities.

## 1 Introduction

The design and implementation of cellular computations using living matter stands as a central goal within the field of synthetic biology [1–4]. Gene regulatory networks are routinely rearranged or entirely synthesized within bacterial [5], yeast [6] and mammalian cells [7], as well as across cellular consortia [8], to manipulate the flow of information from inputs to outputs. This manipulation is achieved by embedding instructions into DNA sequences, specifying which inputs to read and dictating subsequent actions. This algorithmic process, combined with the characteristics of inputs and outputs, gives rise to living computers that implement a certain theoretical model of computation [9].

Models of computation are typically defined by the types of inputs and outputs they permit, as well as the operations they can employ to transform information. In genetic circuits, information is commonly thought of as encoded in the levels of protein expression, with high and low expressions separated by a threshold. This threshold discretizes the genetic signal into distinct high and low values. This analogy mirrors digital electronic circuits, where a signal voltage is assumed to be deterministically either high or low at any given time, carrying precisely 1 bit of information. The application of this abstraction to molecular signaling is the basis for engineering Boolean logic gates and circuits [10–12], enabling synthetic genetic networks implementing counters [13], adders [14], reconfigurable systems [15, 16], distributed computations [17–19], and much more.

While this abstraction is intuitive and practical, it has its limitations. Molecular signals have evolved to carry information over millions of years, without paying any particular attention to the operation of their electronic counterparts. A notable distinction is that, unlike the voltage across electronic circuits, which can be set at high or low steady values constant in time, molecular signals can fluctuate significantly. These fluctuations, known as gene expression noise [20], encode information that goes unused if we merely abstract a signal as a high or low steady value [21–23]. This inherent variability introduces challenges in achieving precise and predictable outcomes in biological computations. The question naturally arises: is digital logic the most appropriate conceptual framework? And, perhaps more crucially, to what extent can we harness these features to implement more powerful computations than achievable through digital logic? Exploring these questions not only challenges our conceptualization of biological computations but also opens avenues for developing novel frameworks that can better accommodate the dynamic and informative nature of molecular signals.

One attempt to transcend Boolean logic involves the implementation of analogue computations in living cells [24–26]. These examples consider the level of proteins over time as an continuous-valued to operate with. Despite representing a definite step forward in harnessing the full potential of molecular information, we argue—a position that underpins this paper—that the core mechanistic details of the gene expression machinery are inherently discrete. Consequently, they may not be ideal for a continuous paradigm like analogue computing, given that a gene is either expressed or not at any given time, irrespective of the final protein expression level. Indeed, whether a transcription factor is bound to the promoter region constitutes a discrete event responsible for the so-called bursting effect[27], which, in turn, drives gene expression in a pulse-like manner[28]. Considering these discrete mechanisms, two frameworks may be more suited than analogue for molecular physical implementations: stochastic and probabilistic computing.

Stochastic computing [29] and probabilistic computing [30, 31] serve as alternatives to conventional computing approaches, but where operations are still based on individual bits of information, i.e., zeros and ones. At the core of both models lies a discrete event, such as a single gene expression burst. The key distinction, simplifying the analysis, is that stochastic computing considers streams of bits—like whether or not a burst occurs in a given period of time—and encodes information in that series of pulses or bit-stream. In contrast, probabilistic computing assigns a probability to the occurrence of a burst, allowing the probabilistic bit, or p-bit, to be either one or zero at any given time.

Computer science has identified specific application domains where these two models are particularly suited. Notably, interesting implementations of this kind of computing have naturally evolved in brain structures like the visual cortex [32]. Furthermore, physical implementations of p-bits are being applied to quantum computing algorithms [33].

The potential of leveraging stochasticity and probability within genetic computations to enter with the goal of cellular supremacy [9], pushing cellular computers into the kinds of application domains where they will outperform conventional ones. This motivates our moving beyond theoretical considerations, to propose specific genetic implementations of synthetic networks that perform functions unattainable by genetic digital logic circuits. The proposed designs provide a foundation for future experimental validation and contribute to the ongoing exploration of advanced cellular computing capabilities.

## 2 Results

We define and provide examples of two approaches aimed at overcoming the limitations imposed by the currently pervasive digital logic conceptualization of molecular information. The first approach involves considering random pulses of signals, while the second involves assessing the probability of signals being high or low. These two approaches enable the implementation of computations that would present significant design challenges as genetic networks within the constraints of digital logic.

### 2.1 Pulse computing with genetic networks

Random pulse computing (RPC) is a stochastic computing paradigm that encodes information with pulses of some signal variable that arrive at random times [29]. A single pulse is a sudden, transient increase in signal intensity followed by a return to a lower, basal intensity. A series of pulses forms a pulse train, which encodes a continuous variable in the frequency of pulse arrivals. In RPC, consecutive pulses occur independently at random times, and the average frequency at which they occur encodes a continuous variable, a real number.

Gene expression is already well known to be a stochastic and bursty process [34]. We propose that it follows naturally to use bursts of gene expression to represent pulse trains, transmitting and transforming information with pulses of expression in gene regulatory networks. This unconventional way of interpreting gene expression levels will open the door to new kinds of computational operations that may lack concise or intuitive implementations as genetic logic circuits.

RPC presents opportunities to concisely express some computations that are difficult to express in other paradigms [35]. This is most often pointed to as an advantage of stochastic computing in general, where operations such as multiplication, which require entire circuits to achieve with binary encoded variables, can be achieved with a single logic gate [36]. By conceptualising genetic circuits in a RPC framework, we utilise these advantages to propose genetic networks for two operations, a pulse matcher and a pulse differencer, that have potential pattern matching applications [37], and which implement computation for which it is hard to even imagine a network of genetic logic gates.

#### 2.1.1 Incoherent feedforward loops as pulse repeaters

A potential advantage of random pulse computing is the robustness it offers with respect to the parameters of the individual genetic components, and the reliability with which pulse encoded signals are propagated through networks.

We investigate this using the simplest operation possible on a pulse train, repeating it. A genetic pulse repeater can be implemented using an incoherent feedforward loop (IFFL), which are the classic pulse generating genetic network motifs [38]. A network diagram of a type 3 IFFL is shown in Figure 1A. This three node network motif responds to pulses of the input signal (*X*) by producing a corresponding pulse of the output (*Z*) at some time later. To this end, the network includes an intermediary node *Y*, which induces expression of the output *Z* unless it is repressed by the input *X*. In the absence of the input, the network remains silent, with no expression of its nodes. However, upon an input pulse, the intermediate signal *Y* is produced. In this state, when the input fades away, the output pulse of *Z* is activated.

**Figure 1:**
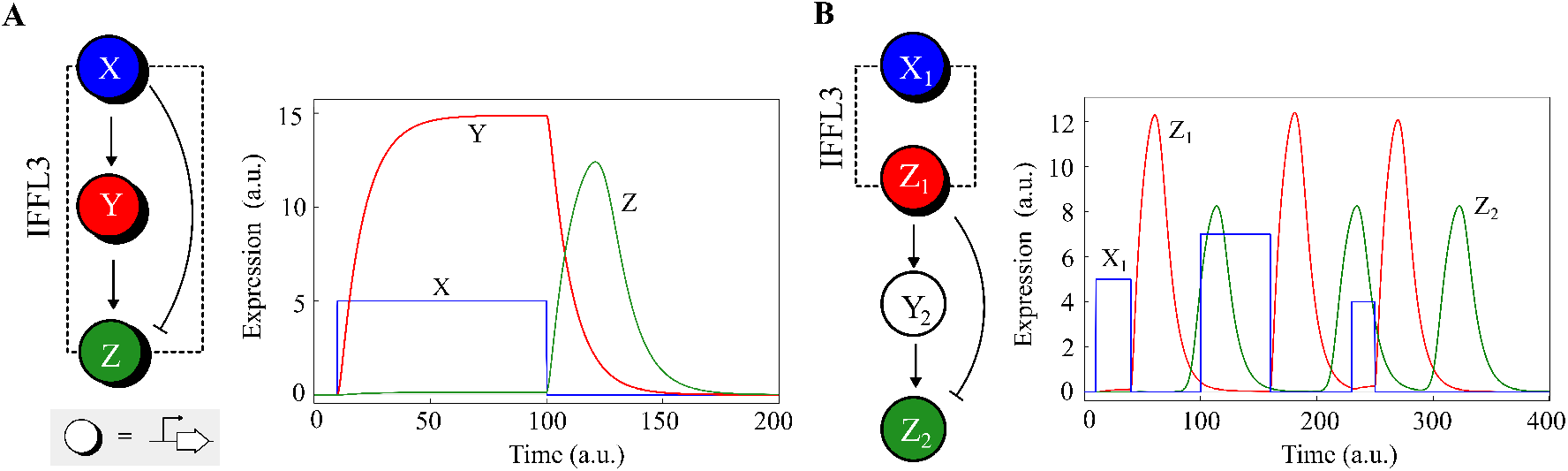
Pulse trains: the propagation of input pulses in gene networks. **A**. The network topology of the type 3 incoherent feed-forward loop (IFFL3). The regulatory node at the input *X* represses the output *Z* but induces expression of an intermediary *Y*. *Y* induces expression of *Z*, but only in the absence of *X*. The plot shows a simulation of the IFFL3 responding to an input pulse of *X* (blue). In the absence of the input the network does nothing, but upon the rising edge of an input pulse, *Y* (red) is expressed, while expression of *Z* (green) is blocked by *X*. Upon the falling edge of the input pulse, an output pulse of *Z* is produced. **B**. IFFLs can be chained together to propagate the pulse train. The output *Z* of the preceding IFFL in the chain in used as the input of the next. The plot shows a simulation of a chain of two *repeaters*. A square pulse (blue) is propagated to the output of a first repeater (red) and then onto a second (green). The pulses do not have to be of a specific amplitude or width, so long as they are sufficient to trigger a pulse in the downstream repeater. Repeaters simply propagate a pulse train, the output of the repeater is the same Poisson point process as at the input.

An IFFL is a pulse repeater and simply propagates frequency encoded information from its input to its output. To do this well the average frequency of the output pulses should be the same as the average frequency of the input pulses. In this regard there are important points to be made about the reliability of the IFFL as a pulse repeater.

Firstly, the magnitude of the delay between an input pulse and the corresponding output pulse does not matter and does not attenuate the output pulse frequency. Furthermore, a stochastic delay can be admitted so long as the individual delay times are independent. This follows directly from the displacement theorem for Poisson point processes, by viewing the delay of a repeater in propagating a pulse as a random displacement. Under random displacement the theorem proves that the output is the same Poisson point process as the input.

Secondly, the IFFLs tend to produce an output pulse with a maximum width which depends on the kinetic parameters of its components, not the width of the input pulse. So a particular pulse repeater will reliably produce a fixed width pulse, in response to input pulses of variable widths. Of course in practice there will be a minimum input pulse width required to produce a response, but from this lower bound upwards, the repeater will be robust to variability in input pulse width. An example is shown in the simulation in Figure 1B, where various different input pulse widths produce identical output pulse responses. In this example, two IFFL motifs are chained together to propagate an input pulse train through both of them, where the output of the first IFFL in the chain is used as the input for the next. The robustness with which repeaters propagate pulse trains constitute an advantage when compared against the variability in signal processing when encoding information in threshold-based digital bits.

#### 2.1.2 Computing with pulses

A common, and useful, approach to decoding the information in a pulse train is to discretise time into bins and estimate the probability of observing a pulse each time bin [35]. This kind of decoding interprets the pulse train as a point process, disregarding the shape, width or amplitude of the pulse itself. In practice of course, pulses are not points and have a finite duration, but it is a common simplifying assumption that we will take advantage of to formalise our computing operations mathematically. Under these assumptions a single genetic pulse train encodes a real number, *p* ∈ [0, 1], which is the probability of finding a pulse of expression in a particular finite time period, and operations transform a collection of probabilities into another collection of probabilities.

The kind of idealised repeater shown in the previous section performs the null operation. It simply takes the input pulse train and copies it at the output, taking *p* as an input and producing *p* as an output. However, a non-idealised repeater is subject to errors. In particular, the repeater might fail to produce an output pulse upon receiving an input pulse, that is, it might “drop” pulses from the input.

We can express the effect of this error mathematically. If we assume a faulty repeater has some rate of error *σ*, then the probability of finding a pulse at the output is the original probability at the input, multiplied by the probability that an error doesn’t occur. So if we assume that the input to a faulty repeater is a pulse train encoding probability *p*, then the output will be another pulse train encoding probability *p*(1 − *σ*). In other words, the output is the input scaled according to the error rate of the repeater, using the noise in the underlying genetic processes for transforming the input. It is promising that we might in the future be able to engineer gene networks whose errors we could actually utilise to perform a mathematical operation.

It is possible to build complex operations using multiple repeaters connected together in larger networks. Figure 2 shows two such networks, the matcher and the differencer.

**Figure 2:**
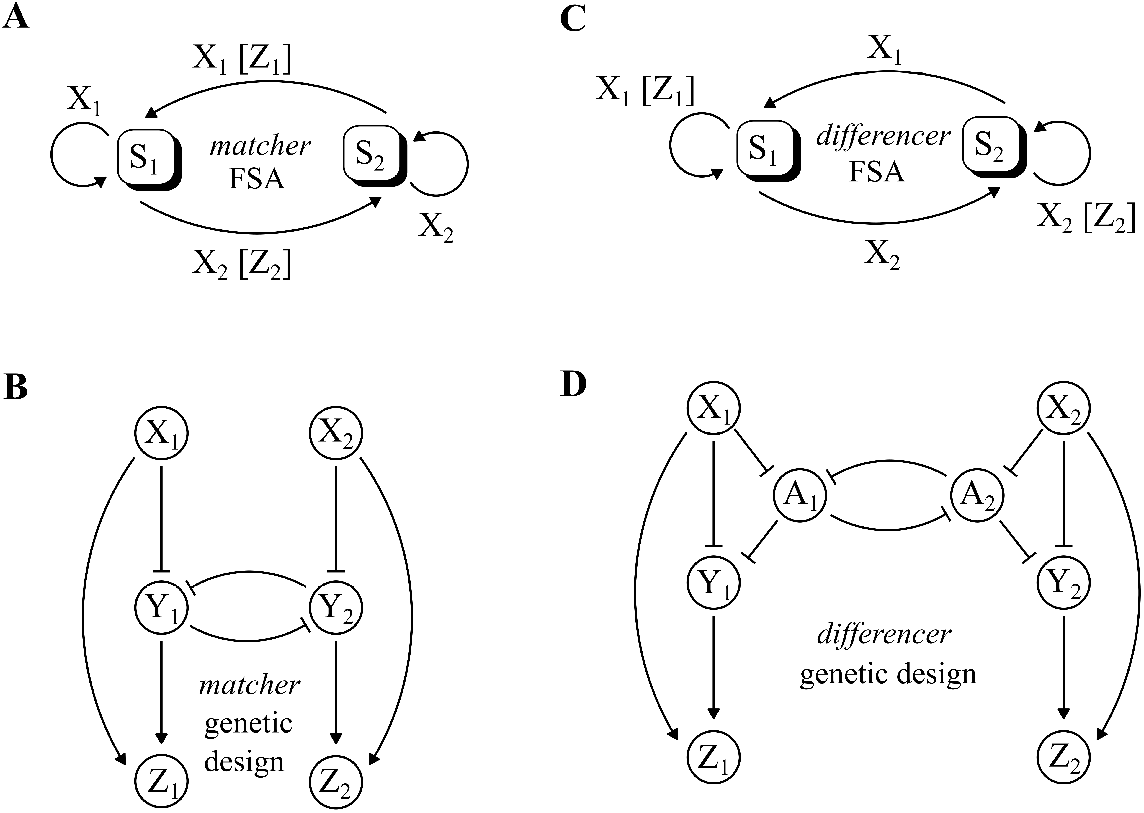
Pulse-based operations: *matcher* and *differencer*. This figure shows two kinds of operation acting on input pulse frequencies, the matcher, and what could be considered its inverse, the differencer. *X*1 and *X*2 are input pulse trains, and *Z*1 and *Z*2 are output pulse trains. **A**. A finite state automata for the matcher operation. There are two states *S*1 and *S*2, switched between by pulses of *X*1 and *X*2. The outputs only pulse upon switches between states; the matcher’s outputs pulse only when pulses of *X*1 are followed (matched) by pulses of *X*2, and vice versa. **B**. A genetic network which implements the matcher by merging two IFFLs with a toggle switch. **C**. A finite state automata for the differencer operation, which is identical to the matcher apart from the transitions where output pulses are emitted. Now the outputs only pulse when the automata stays in the same state; the differencer’s outputs pulse only when there are two consecutive pulses of *X*1 or *X*2. **D**. A genetic network which implements the matcher, using *A*1 and *A*2 as additional nodes which join together two IFFLs.

The operation of the matcher is shown in Figure 2A as a finite state machine with two states *S*_1_ and *S*_2_. While in *S*_1_ the matcher waits for a pulse of the input *X*_2_ to appear. When it receives *X*_2_, the matcher moves to state *S*_2_ and outputs a pulse of *Z*_2_. While in *S*_2_ the matcher waits for *X*_1_, before moving to *S*_1_ and outputting *Z*_1_. This machine can be implemented in a genetic network by coupling two repeaters with a mutual inhibition between their respective *Y* nodes, as shown in Figure 2B. This network has two inputs and two outputs, and the outputs only pulse when the input pulses alternate (a pulse of one input is “matched” by a pulse of the other input). A simulation of the network in Figure 3A shows an example. When a pulse of *X*_1_ arrives directly after a pulse *X*_2_, then a pulse of *Z*_1_ is produced. Similarly a pulse of *X*_2_ after a pulse of *X*_1_ produces a pulse of *Z*_2_. However, two consecutive pulses of either *X*_1_ or *X*_2_ do not produce any output pulses.

**Figure 3:**
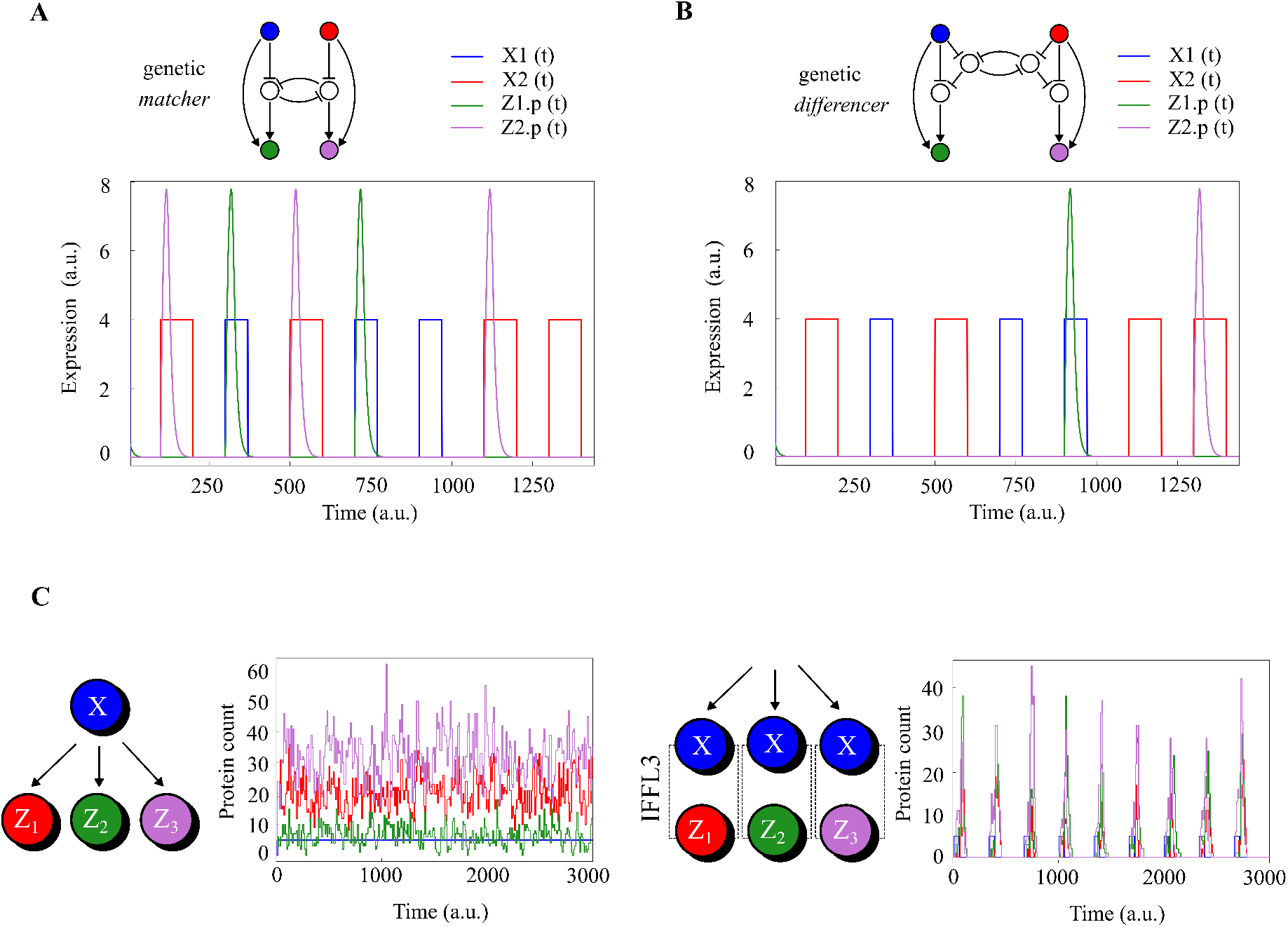
Pulse-based computations: performance and robustness. **A**. This figure shows a time course simulation of the matcher network in operation. *X*1 pulses (blue) followed directly by *X*2 pulses (red) produce *Z*2 pulses (purple), and *X*2 pulses followed directly by *X*1 pulses produce *Z*1 pulses (green). Two consecutive *X*1 or *X*2 pulses produce no response in *Z*1 or *Z*2. The matcher’s outputs have a frequency that is the minimum frequency of the inputs. **B**. This figure shows a time course simulation of the differencer network in operation. Two consecutive *X*1 (blue) or *X*2 (orange) pulses produce pulses in *Z*1 (green) or *Z*2 (purple) respectively. *X*1 pulses followed directly by *X*2 pulses, or vice versa produce no response in *Z*1 or *Z*2. The frequency of the outputs of the differencer reflect the magnitude and sign of the differencer between the two input frequencies. **C**. Demonstration of the robustness of the pulse train propagation as compared to propagation by straightforward genetic regulation. On the left a single transcriptional activator *X* controls expression of three different genes *Z*1, *Z*2 and *Z*3. If expression from these downstream genes has different parameters, then the input signal *X* also propagates differently to output *Z* gene expression. This can be seen in the stochastic simulation. In contrast, an equivalent genetic network which propagates a pulse train encoded signal is shown on the right. Pulse arrival rates do not change with the parameters of the network. This is shown in the simulation, where each IFFL has different transcription rates, but their average output pulse arrival rate is unaffected.

The output *Z*_1_ only pulses when there is a pulse of *X*_1_. However, there is an additional condition that the previous pulse must have been *X*_2_. So if we let *p*_*X*1_ and *p*_*X*2_ be the probabilities corresponding to the inputs *X*_1_ and *X*_2_ respectively, then we can calculate the probability *p*_*Z*1_ of the output *Z*_1_ as:

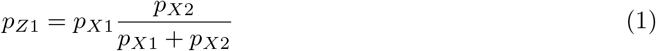

Where 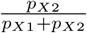 is the probability that the previous input pulse was *X*_2_ instead of *X*_1_. Applying the same to the output *Z*_2_ shows that actually the two outputs of the matcher encode the same probabilities.

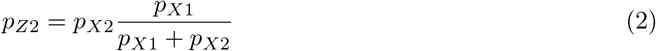

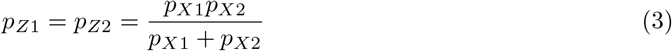

Written this way, matching operation is an approximation to a kind of scaled minimum function of the two input probabilities. In other words, an output pulse of the matcher requires a pulse from both inputs, so that the probability of an output cannot be any greater than the probability at the least likely input. Another way of thinking about this, as an operation on pulse trains, is that the matcher takes two input pulse trains and produces two output pulse trains that are more similar (“matched”) than the inputs.

The differencer can be viewed as an opposite of the matcher, and is shown as a finite state machine in Figure 2C. It has the same states and transitions as the matcher, but produces output only when the machine stays in the same state after receiving an input. The differencer‘s two outputs *Z*_1_ and *Z*_2_ respectively count consecutive pulses of the two inputs *X*_1_ and *X*_2_. In our design for the genetic network implementing the differencer, we couple two repeaters using two additional nodes *A*_1_ and *A*_2_, as shown in Figure2D. A simulation of this network in Figure 3B shows how the outputs *Z*_1_ and *Z*_2_ only respond to consecutive *X*_1_ or *X*_2_ pulses, and a comparison with the simulation of the matcher in Figure3A illustrates the sense in which the difference operation is the inverse of the match operation, in that pulses in the matcher‘s outputs occur only where there are no pulses in the differencer‘s outputs, and vice versa.

For the differencer, the output *Z*_1_ only pulses when there is a pulse of *X*_1_, but this time the additional condition is that the previous pulse must have been *X*_1_ also. For input probabilities *p*_*X*1_ and *p*_*X*2_ we have that:

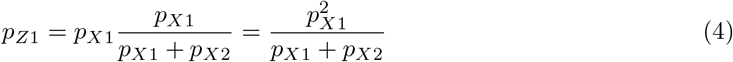

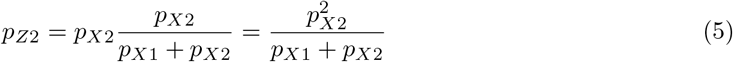

In constrast to the matcher, the differencer produces two different probability outputs that amplify the differences between the inputs. In terms of pulse trains, and again in contrast to the matcher, the differencer takes two input pulse trains and produces two output pulse trains that are more different than the inputs.

#### 2.1.3 Promise of robustness and modularity

In Figure 1B we show that IFFLs can be chained together and reliably propagate the information encoded by a pulse train, even if the shape, width and amplitude of the pulses produced by each IFFL are different. We can hypothesise that this robustness will make genetic pulse computing more forgiving than other paradigms, for example genetic logic computing, in terms of the parameters of components in the networks.

Compatibility of genetic components is one of the principle concerns in the synthesis of genetic logic circuits. The problem arises because differences in the parameters of the components lead to different interpretations of which expression levels correspond to a logic 0 or 1 [39]. In the case of genetic pulse computing, the arrival rate of the output pulse trains does not change depending on the parameters of the individual networks, and the frequency encoded signal is propagated identically across a broader range of parameters, even in a stochastic setting.

An illustration of this idea is shown in Figure 3C, with simulations of a signal *X* being sensed and propagated by three different genetic networks. On the left, *X* is propagated by networks implementing simple genetic regulation (*X* activates expression of *Z*_1_, *Z*_2_ and *Z*_3_). In this case, the output expression levels of the three networks will depend on the network parameters, that is, the specific genetic components that are used to implement the network. In the context of genetic logic computing, this leads to the problem of “matching” the output expression levels of one genetic logic gate to the input “thresholds” of another, in order to join them together and construct genetic logic circuits.

However, on the right of Figure 3C, a frequency encoded signal *X* is propagated by three different genetic IFFL networks *Z*1, *Z*2 and *Z*3 (pulse repeaters as presented in Section 2.1). The amplitude and shape of the output pulses are different, but the arrival rate is not affected, meaning any reasonable pulsing network can be compatible with any other. This mitigates the problem of having to match up the right components to build more complex circuits, and is also a strategy used in natural biological networks [40].

### 2.2 Probabilistic computing with genetic networks

This section introduces the concept of probabilistic bits (p-bits) through genetic circuits in living cells, as demonstrated in three key models. In essence, a p-bit is defined by the transcriptional activity of a gene: a value of 1 when a gene is transcribed and 0 when it is not. Our models explore the dynamics of gene regulation by transcription factors (TFs) to control these p-bit states. The first model outlines the basic mechanism of a single genetic p-bit. The second and third models extend this concept to construct invertible genetic logic gates (NOT and AND gates), demonstrating how combinations of genes can perform logical operations based on their transcriptional states, effectively translating genetic activity into probabilistic computing.

#### 2.2.1 Conceptualisation of probabilistic bits in living cells

We have developed a new model of p-bits using genes as stochastic units (Figure 4). When a gene is being transcribed by RNA polymerases (RNAp), the p-bit value is assigned as 1. On the contrary, when the gene is not being transcribed, the p-bit value is designated as 0 (Figure 4A). The transition between the two values 0 and 1 is controlled by TFs (Figure 4B), which would allow for RNAp to transcribe the downstream coding sequence. In our model, there are four possible states for a given gene that are indicated with *P, Pa, Pr* and *Pra. P* indicates that there is no TF bound to the promoter region and nor RNAp transcribing the gene. *Pa* denotes the TF bound to the promoter but transcription does not occur. *Pr* depicts transcription by RNA polymerase without the TF bound to the promoter. *Pra* illustrates transcription with the TF bound to the promoter. The states *P* and *Pa* represent the p-bit value 0 (no transcription), while *Pr* and *Pra* represent the p-bit value 1 (transcription). The percentage of transcription time, which is the probability of value 1 for the p-bit, is a sigmoidal function of the production rate of the TF (Figure 4C). Note that a similar graph can be obtained when the variable is the total number of TFs instead of the production rate. In this way, the number of TFs in the cell can control the probability of the two values of the genetic p-bit by a sigmoidal function. The production rate of the TF is *k*_*in*_. If *k*_*in*_ = 0.1 the probability of the value 0 of the p-bit is higher than value 1, if *k*_*in*_ = 4 the probability of the values 0 and 1 are similar and if *k*_*in*_ = 10 the probability of the value 1 of the p-bit is higher than value 0 (Figure 4D).

**Figure 4:**
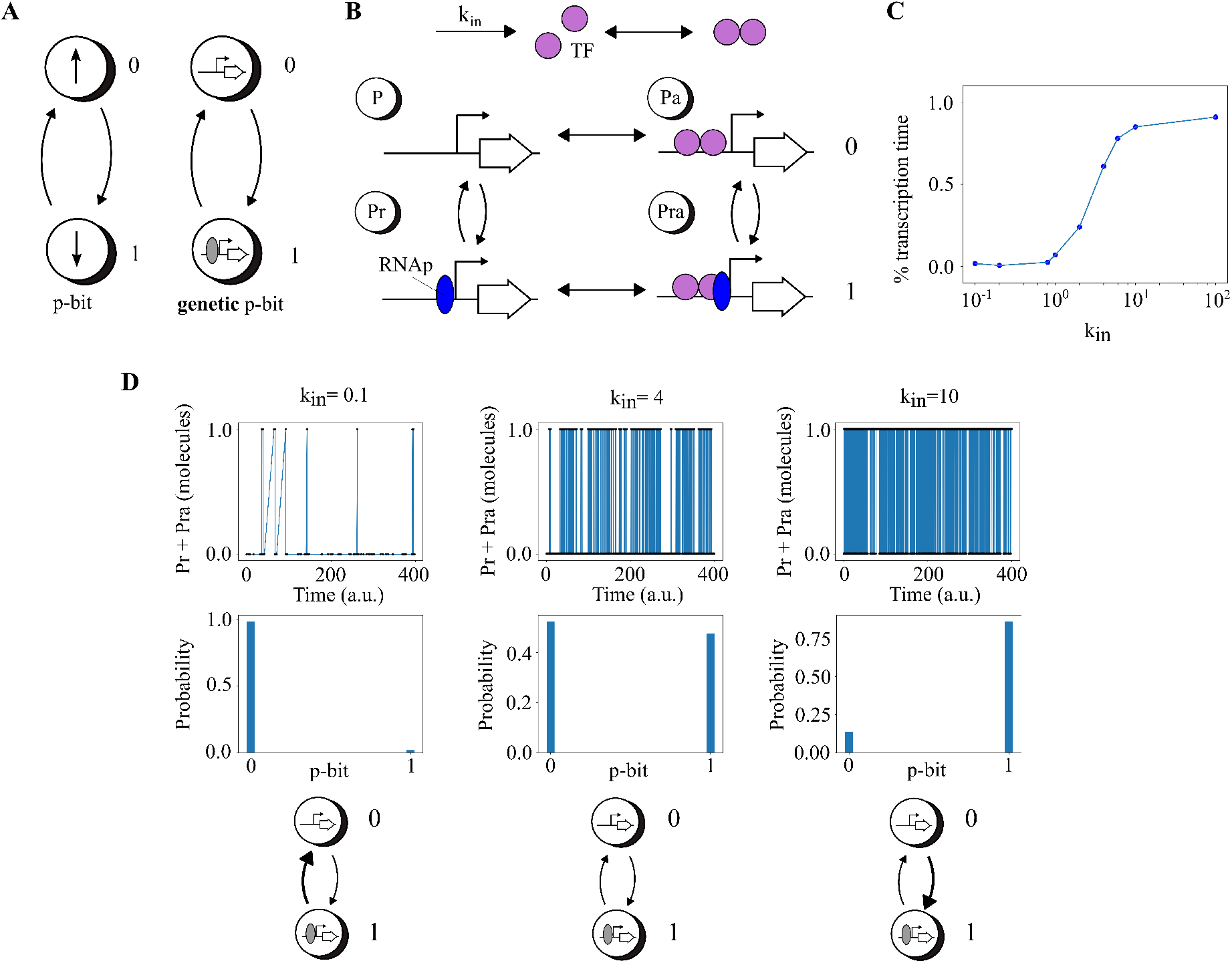
A p-bit using genetic networks in living cells. **A**. General concept of a p-bit and our implementation of a genetic p-bit. When the gene is being transcribed by RNA polymerases, the p-bit value is assigned as 1. Conversely, when the gene is not being transcribed, the p-bit value is designated as 0. **B**. Detailed model of the genetic p-bit when the gene is regulated by a TF. In this model, there are four states for the gene. *P* indicates that there is no TF and there is no RNA polymerase. *Pa* denotes the TF bound to the promoter without transcription. *Pr* depicts transcription by RNA polymerase. *Pra* illustrates transcription with the TF bound to the promoter. The states *P* and *Pa* represent the p-bit value 0 (no transcription), while *Pr* and *Pra* represent the p-bit value 1 (transcription), respectively. *k*_*in*_ is the production rate of the TF. **C**. Transcription time as a function of *k*_*in*_ is a sigmoidal function. **D**. Behavior of the model for three representative values of *k*_*in*_ with their probabilities.

#### 2.2.2 P-bit invertible logic gates

Following the above definition of a genetic p-bit, we now describe the potential implementations of logic gates with *inversion* capabilities. The advantage of using p-bits its that the logic gates can work in two different modes: direct and inverted. The direct mode is the regular one in which the inputs are pinned in order to obtain the output. In the inverted mode, the output is pinned and the inputs fluctuate in order to give all the inputs at once compatible with that output. In order to demonstrate this invertible logic operation we have implemented two models of logic gates with genes: a NOT gate and an AND gate.

The invertible NOT gate (Figure 5A) is a two p-bits network involving two genes that mutually repress each other. Specifically, the protein expressed by gene 1, a TF, binds to the promoter of gene 2 and inhibits its transcription (Figure 5B). Conversely, gene 2 expresses a protein, also a TF, which binds to the promoter of gene 1, inhibiting its transcription. Here, the input is a p-bit implemented by gene 1, and the output is a p-bit implemented by gene 2. When a gene is being transcribed, the p-bit value is 1; when it is not, the value is 0. In direct mode, the NOT gate’s input is fixed (clamped) to 1 or 0, while the output fluctuates between 0 and 1 with varying probabilities (Figure 5C and D). If the input is clamped to 1, the output is more likely to be 0 than 1. Conversely, if the input is clamped to 0, the output is more likely to be 1 than 0, embodying the NOT gate logic. In inverted mode, the NOT gate’s output is clamped, and the input fluctuates, reversing the logic (Figure 5E and F). Here, if the output is clamped to 0, the input is more likely to be 1 than 0, and vice versa. This configuration allows for deducing the input from the NOT gate’s output.

**Figure 5:**
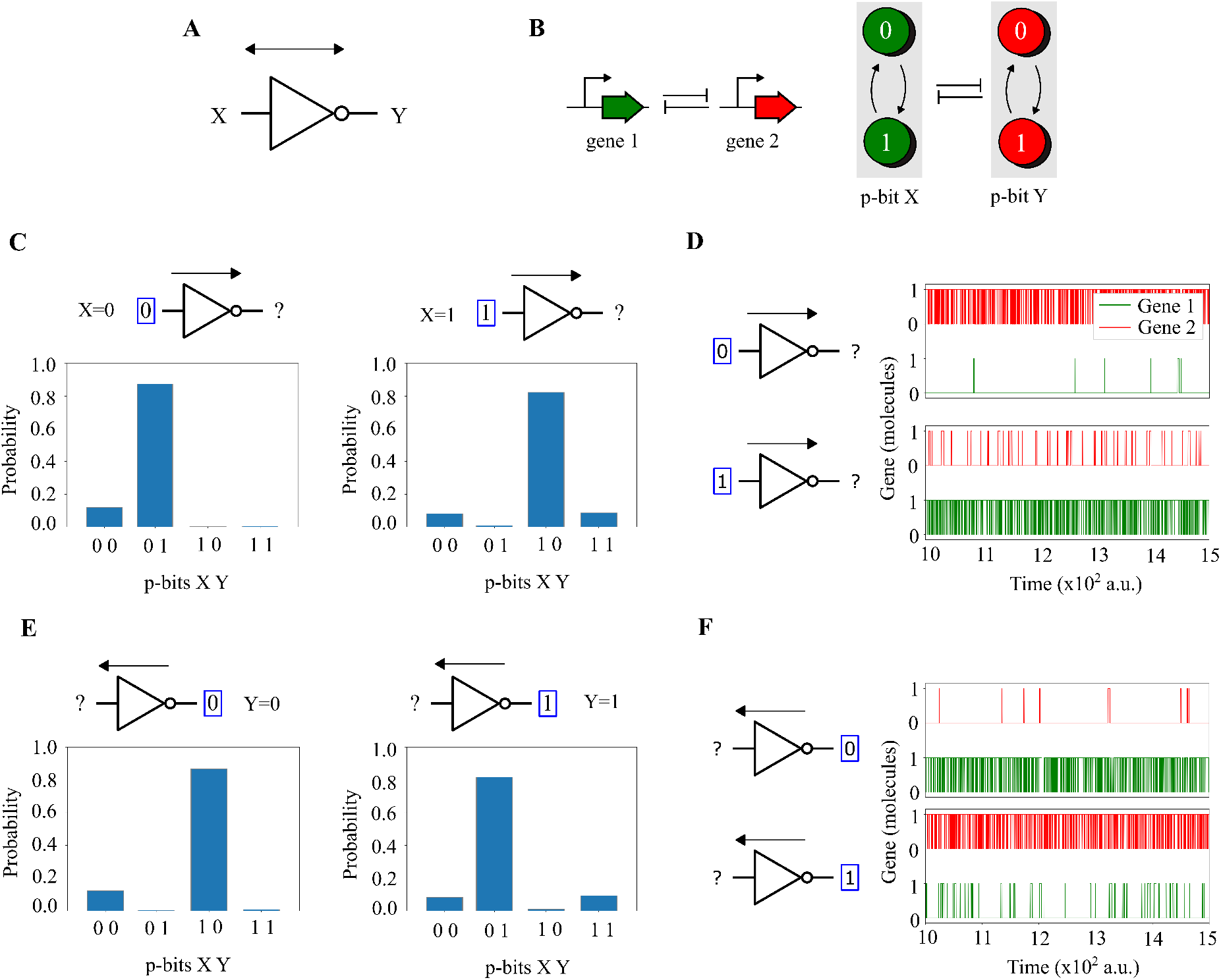
Two-p-bits NOT gate using genetic networks. **A**. Invertible p-bit NOT gate with input *X* and output *Y*. **B**. Invertible p-bit NOT gate implemented with genes. The network consists of two genes mutually inhibiting each other’s transcription, functioning as input (gene 1) and output (gene 2) p-bits. Transcription of each gene sets the corresponding p-bit to 1, while non-transcription sets it to 0. *X* and *Y* represent the p-bits of gene 1 and gene 2, respectively. **C**. and **E**. Invertible p-bit NOT gate operation. The gate operates in two modes: direct, where input is fixed and output fluctuates, and inverted, where output is fixed and input fluctuates. In direct mode (**C**), the NOT gate’s input is set to 1 or 0, causing the output to vary between 0 and 1. A clamped input of 1 results in an output of 0, and vice versa, demonstrating NOT gate logic. In inverted mode (**E**), the output of the NOT gate is set to a constant value, either 0 or 1, which then influences the state of the input. When the output is fixed at 0, the input tends to be 1, and when the output is fixed at 1, the input tends to be 0, enabling deduction of the input based on the output. **D**. Time Evolution in Direct Mode: With the input at 0, only Gene 2 is transcribed, while Gene 1 remains inactive. Conversely, setting the input to 1 activates Gene 1 and deactivates Gene 2. **F**. Time Evolution in Inverted Mode: Fixing the output to 0 activates Gene 1 and deactivates Gene 2. With the output at 1, the situation reverses, activating Gene 2 and deactivating Gene 1.

The invertible AND gate (Figure 6A) is a network of three p-bits involving three genes. Genes 1 and 2 are mutually repressed, similar to the NOT gate, while genes 1 and 3, and genes 2 and 3, mutually activate each other (Figure 6B). The proteins expressed by genes 1 and 2, which are TFs, bind to the promoter of gene 3, increasing its transcription rate. Conversely, the protein expressed by gene 3, also a TF, binds to the promoters of genes 1 and 2, enhancing their transcription rates as well. In direct mode, the two inputs are clamped to 00, 01, 10, or 11, and the output is free to fluctuate between 0 and 1 with different probabilities for each value (Figure 6C and D). If the two inputs are clamped to 00, 01, or 10, the probability of a 0 in the output is higher than that of a 1. However, if the two inputs are clamped to 11, the probability of a 1 in the output is higher. This configuration aligns with the logic of an AND gate. In inverted mode, the output of the AND gate is clamped to 1 or 0, and each input can fluctuate between 0 and 1 (Figure 6E and F). In this scenario, if the output is clamped to 0, the probabilities of 00, 01, and 10 in the two inputs are higher than that of 11. Conversely, if the output is clamped to 1, the probability of 11 in the two inputs is higher than those of 00, 01, and 10. This setup allows us to deduce the inputs from the output of the AND gate.

**Figure 6:**
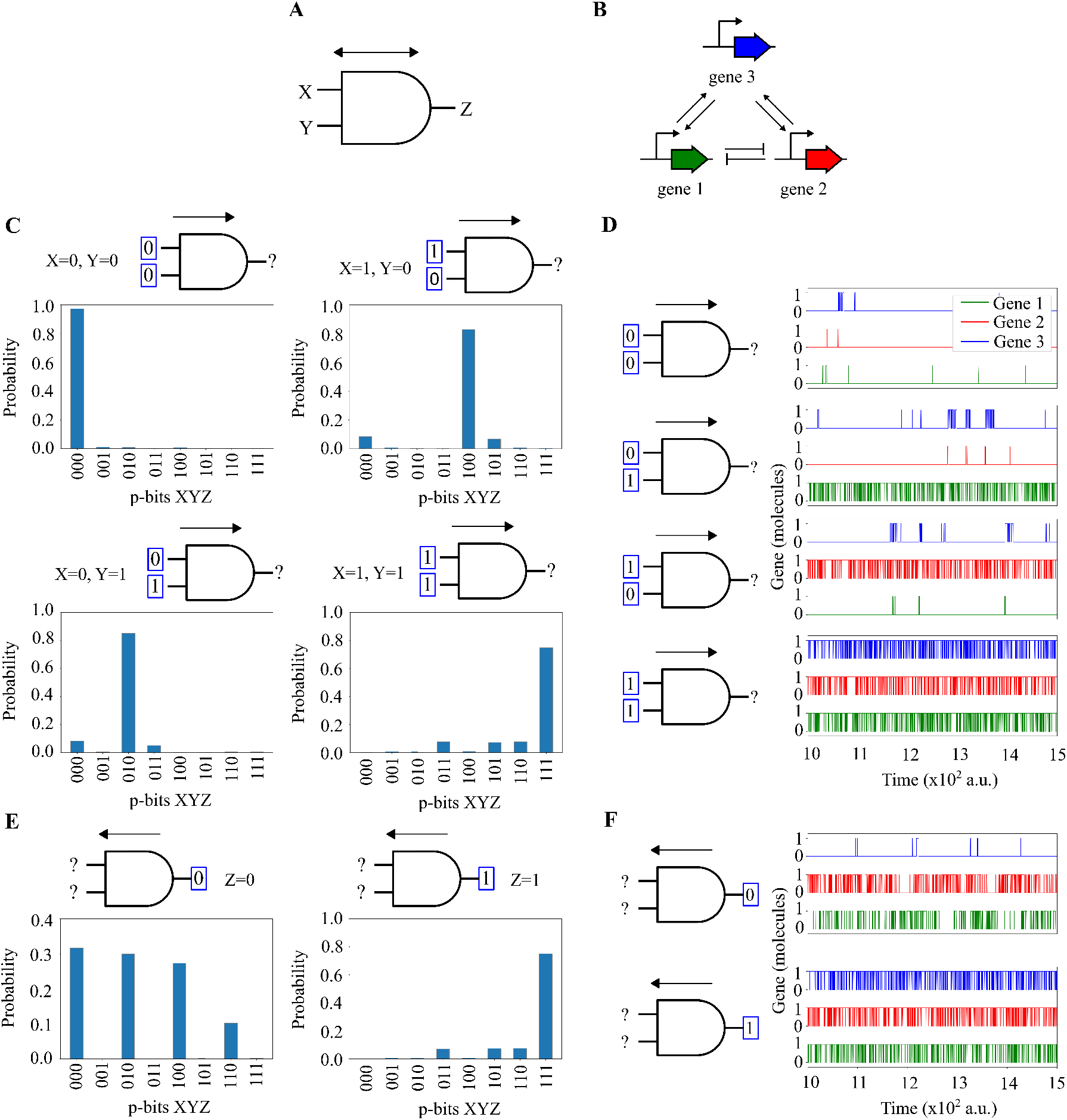
Three-p-bits AND gate using genetic networks. **A**. Invertible p-bit AND gate with inputs X,Y and output Z. **B** Invertible p-bit AND gate implemented with genes. This network involves three genes, with genes 1 and 2 mutually repressing each other and genes 1 and 3, and 2 and 3, mutually activating each other. TFs from genes 1 and 2 increase gene 3’s transcription, while gene 3’s factor enhances genes 1 and 2. *X, Y* and *Z* represent the p-bits of genes 1, 2 and 3, respectively. **C**. and **E**. Invertible p-bit AND gate operation. In direct mode (**C**), inputs are set to 00, 01, 10, or 11, influencing the output’s fluctuation between 0 and 1. A 0 output is more likely for inputs 00, 01, or 10, while a 1 output is likelier for input 11, replicating AND gate logic. In inverted mode (**E**), a fixed output (0 or 1) affects the probabilities of input combinations. With output set to 0, inputs 00, 01, or 10 are more probable, and with output at 1, input 11 is likelier, enabling deduction of inputs from the output. (Continued on the following page.) **D**. and **F**. Time Evolution of the AND Gate in direct and inverted modes. In direct mode (**D**), the transcriptional activity varies based on input combinations. With input 00, none of the genes are transcribed. For input 01, only Gene 1 is transcribed as Genes 2 and 3 remain inactive. Input 10 leads to the transcription of Gene 2 while Genes 1 and 3 stay inactive. With input 11, all three genes are transcribed, reflecting the logical operation of an AND gate. In the inverted mode (**F**), the transcription of genes adjusts based on the output state. With the output fixed at 0, Genes 1 and 2 are transcribed, leaving Gene 3 inactive. Conversely, setting the output to 1 triggers the transcription of all three genes.

## 3 Materials and Methods

### 3.1 Modeling and simulation of IFFLs

Models of gene expression in the IFFLs include the processes of gene transcription, mRNA translation, mRNA degradation, protein degradation. Arrows in diagrams of genetic networks represent transcriptional regulations, which are modeled by hill equations for activation (pointed arrow) or inhibition (blunted arrow).

For the case of deterministic simulations as in Figures 1, 3A and 3B, the model was transformed into a system of coupled ordinary differential equations and solved using the Tsit5 method from DifferentialEquations.jl [41].

For the case of the stochastic simulations in Figure 3C, the model was transformed into a discrete stochastic model and simulated using the implementation of Gillespie’s direct method, also accessed through DifferentialEquations.jl [41].

In the interest of reproducability, the code used to generate the models and produce the simulations of the RPC circuits shown in the Figures is made publicly available at:

github.com/BiocomputationLab/genetic-designs-for-stochastic-and-probabilistic-biocomputing.

### 3.2 Modeling and simulation of P-bits

The simulations presented in Figures 4, 5, and 6 were conducted using the Gillespie algorithm [42]. Specifically, the model accounts for transcription, translation, the binding of transcription factors (TFs) to the promoter, detachment of TFs from the promoter, TF dimerization, TF dissociation, and the degradation of mRNAs and TFs. The model for the NOT gate, shown in Figure 5, consists of 30 chemical reactions. Similarly, the model for the AND gate, detailed in Figure 6, includes 99 chemical reactions. Standard values for the reaction rates were used [43]. The code for the NOT and AND gate simulations is publicly available at:

github.com/BiocomputationLab/genetic-designs-for-stochastic-and-probabilistic-biocomputing.

## 4 Discussion

Understanding and utilizing how information is naturally encoded into molecular signals presents a challenge for biocomputing. Imposing a digital logic abstraction onto this process enables the design and implementation of Boolean logic functions within gene regulatory networks, allowing intuitive reasoning about the resulting computations at small scales. For instance, under this paradigm, if a repressor inhibits the expression of a gene, there will simply be no output. While this high-level view has driven success in synthetic biology and biocomputing, a closer examination of the fundamental differences between living substrates and conventional computing technology reveals areas where the abstraction falls short.

For example, the intrinsic stochasticity of molecular interactions bears little resemblance to the transmission of signals across modern digital electronic circuits. Consequently, while averaging the value of a specific molecular signal across a cell population may create the illusion of digital functionality, measurements on a single cell often reveal seemingly random, if not entirely chaotic, dynamic behaviours. Adding to this complexity, cells undergo evolution, an open-ended process where cells continually modify their hardware and software in an ongoing quest for survival and efficiency. These modifications undoubtedly impact molecular signals, adjusting their performance on a much more complex scale than simple binary values of 0 and 1.

Here, we propose utilizing the dynamics that typically cause the digital logic abstraction to fail, turning them into advantages instead. We draw upon computing paradigms that encode information using noise and time, exemplified by two paradigms that surpass the capabilities of combinatorial logic: random pulse computing and probabilistic computing, and suggest genetic implementations of both.

The idea of a random pulse computing (RPC) as a stochastic computing paradigm has been around since at least the year 1956 [29] and its advantages in terms of robustness well known. Unfortunately for RPC, conventional computing was able to solve electrical engineering challenges that offered the same robustness and easier abstractions in the form of digital logic circuits. However, it is clear now that solving these same engineering challenges in living systems will be much harder than in electronics (if at all possible), and so RPC may yet emerge as a prominent paradigm for biocomputing.

Of course, the implementation of RPC with genetic circuits, as we propose here, is not without significant challenges. Notably, the theory of automated RPC circuit synthesis is less well developed than its digital circuit counterpart, and rely on heuristics in order to search the solution space [34, 44]. But there are also more technical questions to be answered about the genetic implementation presented here, especially as they relate to the assumptions taken about the duration, overlap and error rates of genetic pulses and the circuit which operate them.

Traditional computers process information in a deterministic way using bits. Although these computers have become highly advanced, they are unable to effectively solve certain complex problems such as invertible logic. The development of quantum computing uses qubits that can exist in both 0 and 1 states simultaneously, potentially providing a more efficient solution. However, quantum computing has issues like the need for extremely cold temperatures for operation. Probabilistic computing is an alternative that shares some ideas with quantum computing but it can operate at room temperature. It represents a paradigm where uncertainty is inherently managed through probabilistic bits [31]. These p-bits serve as the fundamental building blocks in probabilistic computing systems, enabling computations that effectively handle stochastic processes. Recent advances have incorporated probabilistic computing into real-world devices, using technologies such as stochastic magnetic tunnel junctions [33] and memristors [45] to bridge these concepts with practical electronic systems. Building on these developments, our study introduces a novel approach: implementing probabilistic computing within living cells. This work extends the application of p-bits from electronic devices to biological systems, potentially revolutionizing biocomputing. Our approach takes advantage of the inherent stochasticity of biological processes to mimic these computational structures. Unlike electronic systems, where noise is often a limiting factor, the stochastic nature of gene expression becomes a beneficial feature in our biological p-bit models, allowing for a more versatile and robust computational framework. The direct and inverted modes of operation demonstrated in the NOT and AND gates underscore the potential for genetic circuits to perform flexible, invertible computing, comparable to traditional logic gates but with the added advantage of biological integration.

While our models serve as an initial demonstration, the practical implementation of genetic p-bits faces challenges, including the efficient design of promoters and transcription factors to achieve desired p-bit states reliably. Future research should focus on enhancing the orthogonality and specificity of these genetic components to enable more complex genetic circuits without crosstalk between different p-bits. Additionally, exploring the integration of genetic p-bits with other biological computing components, such as RNA or protein pathways, could further expand the computational capabilities and efficiency of bio-computational systems.

## Author Contributions

All authors conceived the study. LG and JM-B developed mathematical models and performed computational simulations. AG-M supervised the research. All authors contributed to the discussion of the results and wrote the manuscript.

